# Myoglobin promotes cardiomyocyte differentiation through oxidatively modulating the Hippo Kinase Pathway

**DOI:** 10.1101/2022.08.31.506050

**Authors:** Krithika Rao, Elizabeth Rochon, Anuradha Singh, Rajaganapathi Jagnnathan, Zishan Peng, Mousumi Moulik, Manling Zhang, Paola Corti, Sruti Shiva

## Abstract

**Background:** While cardiomyocytes undergo terminal differentiation postnatally and rarely re-enter the cell cycle, the endogenous mechanisms that propagate differentiation and prevent de-differentiation remain unclear. The monomeric heme protein myoglobin, which stores oxygen and regulates reactive oxygen/nitrogen species balance in the heart, increases in expression by over 50% during cardiomyocyte differentiation. Though myoglobin deletion without significant compensation is embryonic lethal in mice, a role for the protein in regulating cardiomyocyte differentiation has not been tested. We hypothesized that myoglobin expression is required for cardiomyocyte differentiation and the loss of myoglobin enables de-differentiation.

**Methods:** Myoglobin was genetically silenced in HL-1, H9C2 cells, and neonatal rat ventricular cardiomyocytes (NRVM) to examine myoglobin-dependent effects on differentiation, proliferation, and Hippo pathway signaling. A zebrafish model of Mb depletion was made using CRISPR-Cas9 to test the effect of myoglobin depletion on cardiac regeneration after apical resection injury in vivo.

**Results:** Myoglobin deletion in cultured cell lines and NRVM decreased the gene expression of cardiomyocyte differentiation markers (*troponin, myosin light chain, and myosin heavy chain*), upregulated markers of dedifferentiation (*runx1* and *dab2*) and stimulated cell proliferation. Mechanistically, we show that the heme prosthetic group of myoglobin catalyzes the oxidation of the Hippo pathway kinase LATS1, which activates the enzyme to phosphorylate the downstream Yes-associated protein (YAP) transcription factor, which prevents its transcriptional activity. Thus, the loss of myoglobin results in the de-phosphorylation and nuclear translocation of YAP, which propagates proliferation and fetal gene expression. *In vivo*, myoglobin-deficient zebrafish hearts recapitulated the increase in YAP signaling and showed accelerated regeneration at 20 days post apical injury.

**Conclusion:** We a novel role for myoglobin as an endogenous driver of cardiomyocyte differentiation, and a regulator of the Hippo pathway. These findings suggest myoglobin as a potential target for strategies to enhance cardiac development and improve cardiac repair and regeneration.

## Introduction

Cardiomyocytes are the primary cell-type responsible for heart muscle contraction. In order to perform this highly specialized function, cardiomyocytes withdraw from the cell cycle soon after birth to become terminally differentiated.^1, 2^ Through the process of differentiation, they acquire the structural, functional and metabolic characteristics necessary to power repeated contraction-relaxation cycles.^3^ While fetal cardiomyocytes are proliferative, differentiated cardiomyocytes rarely re-enter the cell cycle (frequency <1%), in part due to their complex, well-organized cytoskeletal architecture.^4, 5^ A key feature of cardiomyocyte differentiation is the downregulation of fetal gene signatures including cell cycle markers, and upregulation of differentiation markers including mature sarcomeric genes such as *troponin, myosin light chain*, and *myosin heavy chain*.^6-8^ Cardiomyocytes remain terminally differentiated over a lifetime and loss of differentiation (dedifferentiation) is associated with cellular remodeling in multiple pathological conditions.^9^ Though uncommon in mammalian hearts, de-differentiation is also associated with increased cardiomyocyte proliferation and myocardial regeneration, a feature more prevalent in organisms such as the zebrafish (*Danio rerio*).^10-15^ While the phenotypic features distinguishing differentiated from immature or de-differentiated cardiomyocytes are well-characterized, the endogenous mechanisms that drive and maintain differentiation or enable de-differentiation are not well-defined.

Myoglobin (Mb) is a monomeric heme protein constitutively expressed at high micromolar (200-300) concentrations in cardiomyocytes.^16^ The classic function of Mb in cardiomyocytes is considered to be oxygen storage and delivery to support mitochondrial respiration during hypoxia.^17, 18^ More recent studies have identified new roles for Mb in regulating tissue nitric oxide (NO) bioavailability and cellular reactive oxygen species (ROS) levels. Similar to the mechanism of oxygen storage and transport, these functions of Mb are dependent on the protein’s heme prosthetic group. For example, oxidation of the ferric (Fe3+) and ferrous (Fe 2+) forms of the heme center of Mb results in ferryl Mb (MbFe4+), which, through its pseudoperoxidase activity, can oxidize biomolecules such as nucleic acid, amino acids and lipids.^19, 20 21^ *In vivo*, Mb autooxidation under conditions of ischemia/reperfusion has been regarded as a significant source of oxidants that contributes to oxidative damage in the heart.^22, 23^ However, the contribution of Mb’s oxidative signaling to physiological cardiomyocyte function is less well defined. Notably, during the developmental period of cardiac differentiation, the cardiac concentration of Mb increases by over 50% compared with the fetal heart.^24^ Further, global Mb deletion causes embryonic lethality, and mice that do survive to adulthood encompass multiple physiological compensatory adaptations for survival.^25-31^ Despite the requirement for Mb and its dramatic increase in expression during cardiac development, the physiological role of Mb in cardiomyocyte differentiation has not been examined.

The Hippo signaling pathway maintains cardiomyocytes in a differentiated state by integrating extracellular cues with the activity of the transcriptional co-factor yes-associated protein (YAP). ^32-34^ YAP activity is propagated by its dephosphorylation and nuclear accumulation which potentiates a pro-proliferative and fetal gene transcription program. While the embryonic heart relies on YAP activity for proliferation, terminally differentiated cardiomyocytes predominantly express phosphorylated YAP which is sequestered in the cytoplasm for degradation. LATS1/2 is the terminal Hippo pathway kinase which phosphorylates YAP, whose genetic or pharmacological inactivation is sufficient to affect cardiomyocyte differentiation^34, 35^. While recent discoveries reveal that signals besides the canonical Hippo pathway can modify LATS1/2 activity, the effect of endogenous Mb and Mb-dependent signaling including the role of oxidants is unknown ^36-38^.

In this study, we hypothesize that Mb regulates cardiomyocyte differentiation. We demonstrate that deletion of endogenous Mb prevents cardiomyocyte differentiation resulting in the persistence of an undifferentiated, pro-proliferative phenotype. Mechanistically, we show that Mb oxidizes LATS1 to increase its kinase activity and drive differentiation by YAP phosphorylation and inactivation. We demonstrate the activation of this pathway *in vivo* using a cardiac apical resection injury model in a novel mutant zebrafish line lacking Mb in which we show that zebrafish lacking Mb have significantly accelerated scar regeneration post-injury. We discuss the potential implications of this Mb-dependent pathway in the context of understanding cardiac development as well as potential therapeutic strategies.

## Methods

### Cell Culture

H9C2 undifferentiated rat myoblasts were purchased from ATCC and maintained in 10% FBS supplemented with antibiotics and L-glutamine. HL-1 cells were obtained from ATCC and cultured in Claycomb medium supplemented with 10% FBS. H9C2 cells were stimulated to differentiate when cells were at 70-80% confluence using 1% serum medium supplemented with 10nM Retinoic acid, and maintained in differentiation media for 5 days as described.^39, 40^

### Myoglobin knockdown

Transient knockdown of myoglobin in H9C2 rat cells was performed using ON-TARGETplus Rat Mb siRNA (L-094211-02-0005, Dharmacon) or control (scrambled) non-targeting siRNAs (D-001810-01-05, Dharmacon; 50nM). Cells were grown to 70% confluency and siRNAs were transfected with Mirus TransIT-X2 (MIR6004, Mirus Bio) following manufacturer instructions in OPTI-MEM medium (31985070, Thermo Fisher). Cells were analyzed 48 or 72h post transfection. Transient knockdown in mouse HL-1 cells was performed with siRNA particles targeted to mouse myoglobin (sc-35994, Santa Cruz) or non-targeting scramble siRNA (sc-37007, Santa Cruz) following the same protocol.

### Proliferation

Cells were plated in 96 well cell culture plates at a density of 5000 cells/well. 48h later, cells were fixed and stained with crystal violet (0.2%w/v, 30 min), resolubilized in 1% SDS, and absorbance was measured at 595 nm.

### PCR

RNA was extracted using RNAeasy spin columns (#74134, Qiagen) according to manufacturer instructions. Equal concentration of mRNA was measured and used as template for first strand DNA synthesis (#1708891, Biorad). The cDNA was used as input in quantitative RT-PCR reactions using PowerUp™ SYBR™ Green Master Mix (A25742, Thermo Fisher). All primers were obtained from integrated DNA Technologies and sequences are 5’-3’. Rat primers: mlcF (AGGCCTTCACAATCATGGAC), mlcR (TCGTTTTTC ACGTTCACTCG), mhcF (GCCTACCTCATGGGACTGAA),mhcR(ACATTCTGC CCTTTGGTGAC),tnnt2F(CCTGCAGGAAAAGTTCAAGC), tnnt2R(GTGCCTGGCAAGACCTAGAG), 18s F(TTGATTAAGTCCCTGCCCTTTGT), 18sR (CGATCCGAGGGCCTAACTA). Mouse primers: tnnt2F(CAGAGGAGGCCAACGTAGAAG), tnnt2R (CTCCATCGGGGATCTTGGGT), Dab2F(GGCAACAGGCTGAACCATTAGT), Dab2R (TTGGTGTCGATTTCAGAGTTTAGAT), Runx1F (CGGCCCTCCCTGAACTCT), Runx1R (TGCCTGCCTGGGATCTGTA).

### Western blot

Cell lysates were prepared using cOmplete™ Lysis-M (4719956001, Roche) with protease and phosphatase inhibitors and subjected to standard electrophoresis and transfer as previously described^41^. Following transfer, membranes were blocked, incubated in primary antibody (overnight, 4C) followed by incubation in species specific secondary antibody (1:10,000 Li-COR; 2h; room temperature) before imaging on an Odessey CLx (Li-COR Biosciences). The primary antibodies include pYAP (1:500, 4911, Cell Signaling Technologies), Total YAP(1:1000, H-9, Santa Cruz), Mb (1:1000, LS-C144978 LS Bio or 1:1000, sc-393020 Santa Cruz), Tubulin (1:1000, CP06, Calbiochem).

### ROS measurement

200,000 cardiomyocytes were incubated in Amplex Red (5 μM; 2 min) for cellular ROS (A22177, ThermoFisher), or MitoSOX Red (5 μM; M36008, ThermoFisher; time) for mitochondrial ROS. The rate of ROS production was calculated by measuring the linear phase of the kinetic fluorescence emission curves (571/585 nm: Amplex red) or (510/580 nm: mitosox) over 30 mins as previously described ^42, 43^.

### LATS1 kinase activity and oxidation

LATS1 kinase activity was measured in assay was performed in recombinant LATS1 (2ng) or LATS1 immunoprecipitated from cell lysates using an antibody specific to LATS1 (4µg; CST:3777S). Immunoprecipitated LATS1 was incubated with YAP peptide (1µg) and ATP (10 µM). After completion of the reaction (30°C; 30 min), the kinase mix was incubated with ADP-Glo reagent (40 min) and luminescence resulting from ADP conversion to ATP was measured following the manufacturer’s instructions (Promega; ADP-Glo™ Kinase Assay, V6930). To measure changes in oxidation, LATS1 kinase was immunoprecipitated from 500 µg of protein lysates and subject to Oxyblot assay (#S7150, Millipore) according to manufacturer’s instructions.

### Cell Immunofluorescence

Cells were cultured on chamber slides (Millipore C7182-1PAK), fixed in 4% formaldehyde (Thermo 50-980-487; 12 min, 20°C) and then blocked with 5% normal donkey serum (Sigma-Aldrich D9663; 30 min) and Triton X-100 (0.01% in PBS). This followed by incubation in primary antibodies to H3P (1:500; 06-570, Sigma), alpha-sarcomeric actinin (1:500; A7811, Sigma), cardiac troponin (1:500; MA5-12960, Thermo Fisher) and total-YAP (1:250, sc-101199, Santa Cruz) overnight at 4°C. Cells were then washed in PBS (3 times for 5 mins) and incubated at room temperature in Alexa Fluor conjugated secondary antibodies (1 hour;1:500; A-21206, R37115, Thermo Scientific). Cells were washed in PBS (3 times for 5 mins) prior to nuclear counterstaining with DAPI in mounting media (Vector BioLabs).

### Neonatal rat cardiomyocyte isolation

As previously described,^44^ hearts from 1–2-day old Sprague Dawley rat pups were excised into cold buffer (NaCl 137mM, KC1 5.36mM, MgSO4– 7H2O 0.81mM, dextrose 5.55mM, KH2PO4 0.44mM, Na2HPO4–7H2O 0.34mM, and HEPES 20mM at pH 7.5). Ventricles were cleared of blood and dissected into 1–2mm pieces to digest out cardiomyocytes using multiple incubations in trypsin (0.04%) and collagenase (0.4mg/ml) at 37°C on a rocking platform. Cardiomyocytes were then purified from the fibroblasts by pre-plating for 90 mins, and were plated on Matrigel or 0.1% gelatin coated culture dishes prior to performing knockdown studies. Cells were cultured in serum free DMEM containing 0.1% Insulin-transferrin-selenium (Thermo Fisher Scientific).^44^

### Zebrafish myocardial amputation model

A 37 base pair (bp) genomic insertion resulted from a non-homologous end joining repair error following injection of a sgRNA targeting the first exon of zebrafish *mb* using CRISP-Cas9 genome editing. Target site GGATCTGGTTCTGAAGTGCTG (pt813) was selected using CHOPCHOP software. The sgRNA was made according to recommended protocol, using the Megashort SP6 Transcription kit (ThermoFisher, AM1330) followed by RNA purification with the Megaclear Kit (ThermoFisher, AM1908). *Cas9* mRNA was synthesized from pCS2-nCas9n using the mMessage mMachine SP6 Transcription kit (ThermoFisher, AM1340). Approximately 2 nl of the RNA mixture of sgRNA (500 pg) and Cas9 mRNA (600 pg) were coinjected at 1 cell stage. Embryos were raised and screened for changes in PCR amplicon size (mutants being 37 bps longer) using primers specific to exon 1 of *mb* (forward primer: 5’-GCACATCCATACATCGCTTG-3’, reverse 5’– GACAATAATCGAGCAGCCTTAGA–3’).

To induce cardiac regeneration, age matched adult wt and *mb-/-* zebrafish were anesthetized for 3-5 mins in ethyl 3-aminobenzoate methanesulfonate salt (0.168 g/dL). A small incision was made on the ventral wall to access the heart and approximately 20% of the heart ventricle was amputated using a pair of sterile fine scissors, following which fish were allowed to recover in water for 30 min and then returned to the tank until the time of assays^45, 46^.

### Histology, immunofluorescence and imaging of in vivo models

Hearts were collected from sedated zebrafish, washed in cold PBS, and fixed in 4% paraformaldehyde. The hearts were then cryopreserved in a sucrose gradient and embedded in Surgipath Cryo-Gel (Leica). Ten to twelve micrometer cryosections were prepared and consecutive sections were used for Acid Fuchsin Orange G (AFOG) staining or immunofluorescence. The injured area of the zebrafish heart was measured and averaged from the 3 sections displaying the largest injured area in each heart. The primary antibodies used for immunostaining were anti-embCMHC (N2.261, 1:50, DSHB), anti-Mef2C (SC-313, 1:500; Santa Cruz Biotechnology), anti-PCNA (8825, 1:1000; Sigma). The secondary antibodies used for immunostaining (1:1000; Thermo Fisher) were goat-anti-rabbit Alexa Fluor 488 (A11008), goat-anti-rabbit Alexa Fluor Cy3 (A10520), goat-anti-mouse Alexa Fluor Cy3 (A10521), and goat-anti-mouse Alexa Fluor 488 (A11001). Slides were stained with DAPI (1:1000; Thermo Fisher), followed by treatment with Vectashield mounting medium (Vector Laboratories). Confocal images were taken with a Zeiss LSM 200 confocal microscope. Two-dimensional projections were generated from z-series (1 μm steps). Mef2C/Pcna double-positive cells were manually counted over three separate consecutive sections, averaged, and normalized to the total area counted. Only hearts that had a clear injury located in the center sections of the heart were considered for each experiment.

### Statistics

All bar graphs depict mean ± SEM. Two groups were compared for statistical significance using a two-tailed Student’s t-test. For comparison of more than 2 groups, ANOVA following application of Bonferroni post-hoc analysis to correct for multiple comparisons was used. Differences were considered statistically significant when p < 0.05.

## Results

### Endogenous myoglobin is required for differentiation

To determine the role of endogenous Mb expression during cardiomyocyte differentiation we transfected undifferentiated rat H9C2 myoblasts with siRNA and achieved 44.99±10.47% knockdown of endogenous Mb expression compared with non-targeting scramble control (scrm) (**Figure 1A**). Upon stimulation with retinoic acid, Mb-containing control cells differentiated to form morphologically characteristic multinucleated cells and expressed markers of structural differentiation including *myosin light chain, beta-myosin heavy chain* and *cardiac troponin-t* (**Figure 1B**)^39, 40^. In contrast, following identical stimulation, Mb-siRNA cells retained a more undifferentiated mononucleated spindle-shape and did not increase transcript levels of structural differentiation markers (**Figure 1B**). Similarly, in a differentiated HL-1 atrial cardiomyocyte cell line, knockdown of Mb by siRNA (**Figure 1C**) decreased the expression of the differentiation markers *myosin light chain, beta-myosin heavy chain* and *cardiac troponin-t* (**Figure 1D**), and increased the expression of Dab2 and Runx1, markers of de-differentiation (**Figure 1D)**^47^. To test whether Mb impacts differentiation of primary cardiomyocytes, we knocked down endogenous Mb using siRNA in isolated rat neonatal ventricular cardiomyocytes (NRVM; **Figure 1E**). Immunofluorescence staining for cardiac troponin T demonstrated that cells lacking Mb were less successful at forming sarcomeric networks compared to control cells (**Figure 1F**), and had decreased expression of the structural markers *myosin light chain, beta-myosin heavy chain* and *cardiac troponin-t* (**Figure 1G**).

**Figure 1:**
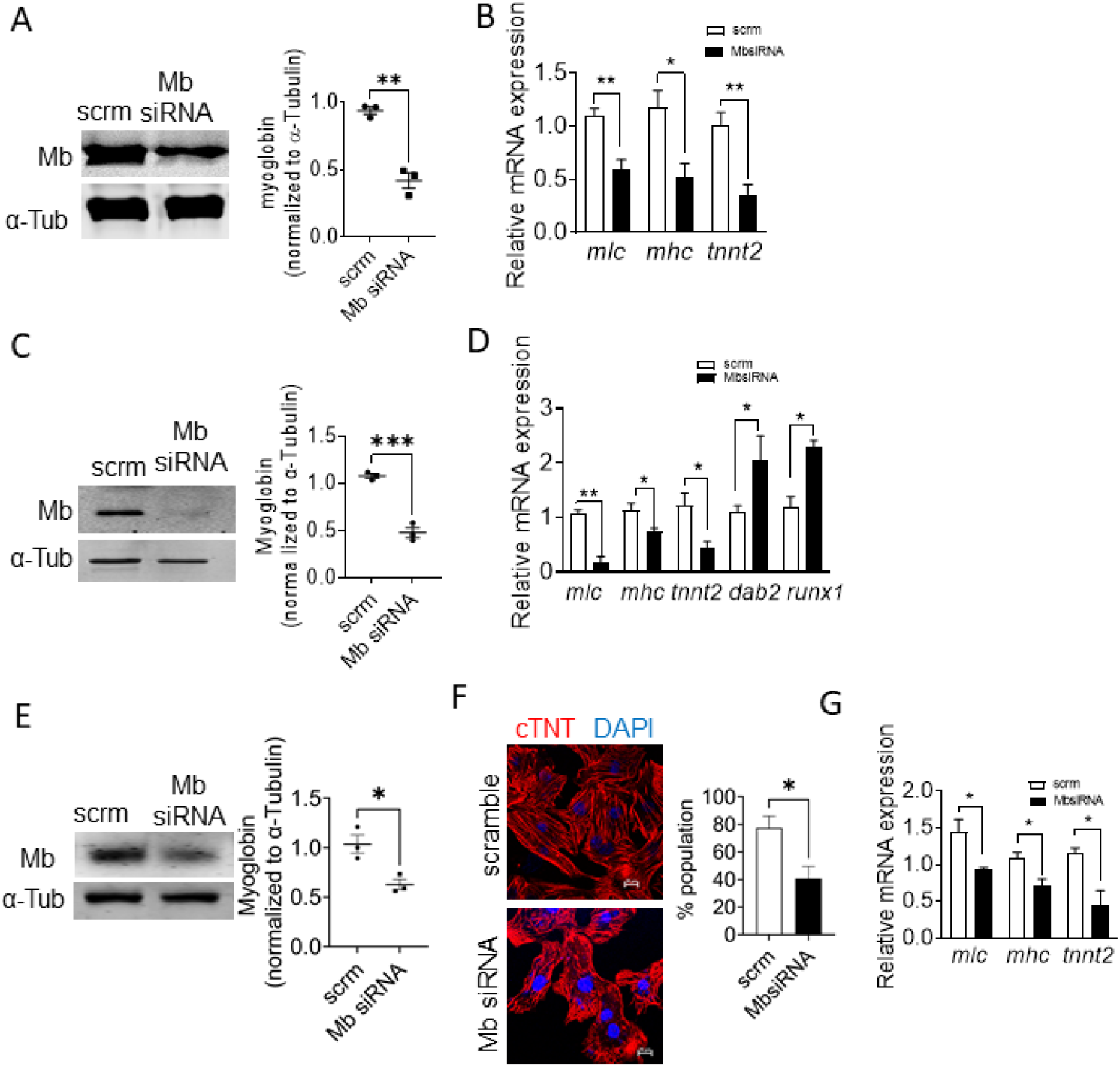
Knockdown of myoglobin inhibits cardiomyocyte differentiation. **(A)** Representative Western blot and quantification of Mb protein in H9C2 rat cardiomyocytes treated with scramble siRNA (scrm) or siRNA targeting Mb (Mb siRNA) and quantification; n=3. **(B)** Levels of cardiac structural differentiation marker mRNA -m*lc (myosin light chain), mhc (myosin heavy chain), tnnt2 (cardiac troponinT*)-in scrm control and siRNA-Mb cells. n=3-4. **(C)** Representative Western blot and the corresponding densitometry for Mb protein in HL-1 mouse atrial cardiomyocytes treated with scrm siRNA or Mb targeted siRNA, n=3. **(D)** Quantification of cardiac structural differentiation markers in HL-1 scrm control and Mb-siRNA cells; n=3. **(E)** Quantification of cardiac dedifferentiation markers in HL-1 scrm control and Mb-siRNA cells; n=4. **(F)** Representative Western blot and quantification of Mb in NRVMs treated with scrm siRNA or Mb targeted siRNA (siRNA-Mb); n=3. **(G)** Representative immunofluorescence images of NRVMs stained for cardiac troponin T (cTNT; red) to visualize sarcomeric structures and DAPI for nuclei (blue). Quantification represents the percentage of NRVMs demonstrating organized, continuous sarcomeres across the cytoskeletal area of the cells; n=200 cells in scrm and Mb siRNA groups. **(H)** mRNA levels of cardiac structural differentiation markers in NRVM after treatment with scrm siRNA or siRNA targeted at Mb. n=3. Bar graphs are mean ± SEM; \**p*< 0.05, \*\**p*< 0.01, \*\*\**p<0*.*001*.

Consistent with an undifferentiated state, H9C2, HL-1, and NVRM cells lacking Mb (Mb siRNA) proliferated at a higher rate compared to control cells expressing endogenous Mb as measured by crystal violet uptake (**Figure 2A-B**) or by immunofluorescence staining for the mitotic marker phospho-histone 3 (H3P) at 48 h post-transfection (**Figure 2C**). Furthermore, compared with control cells, H9C2 Mb-siRNA cells expressed higher levels of the cell cycle drivers Cyclin E and Cyclin A and lower levels of the cell cycle inhibitor retinoblastoma protein (pRb) (**Figure 2D**). Similarly, knockdown of Mb significantly increased NRVM proliferation as detected by H3P staining (**Figure 2E**). Taken together, these data demonstrate that endogenous Mb promotes and maintains cardiomyocyte differentiation.

**Figure 2:**
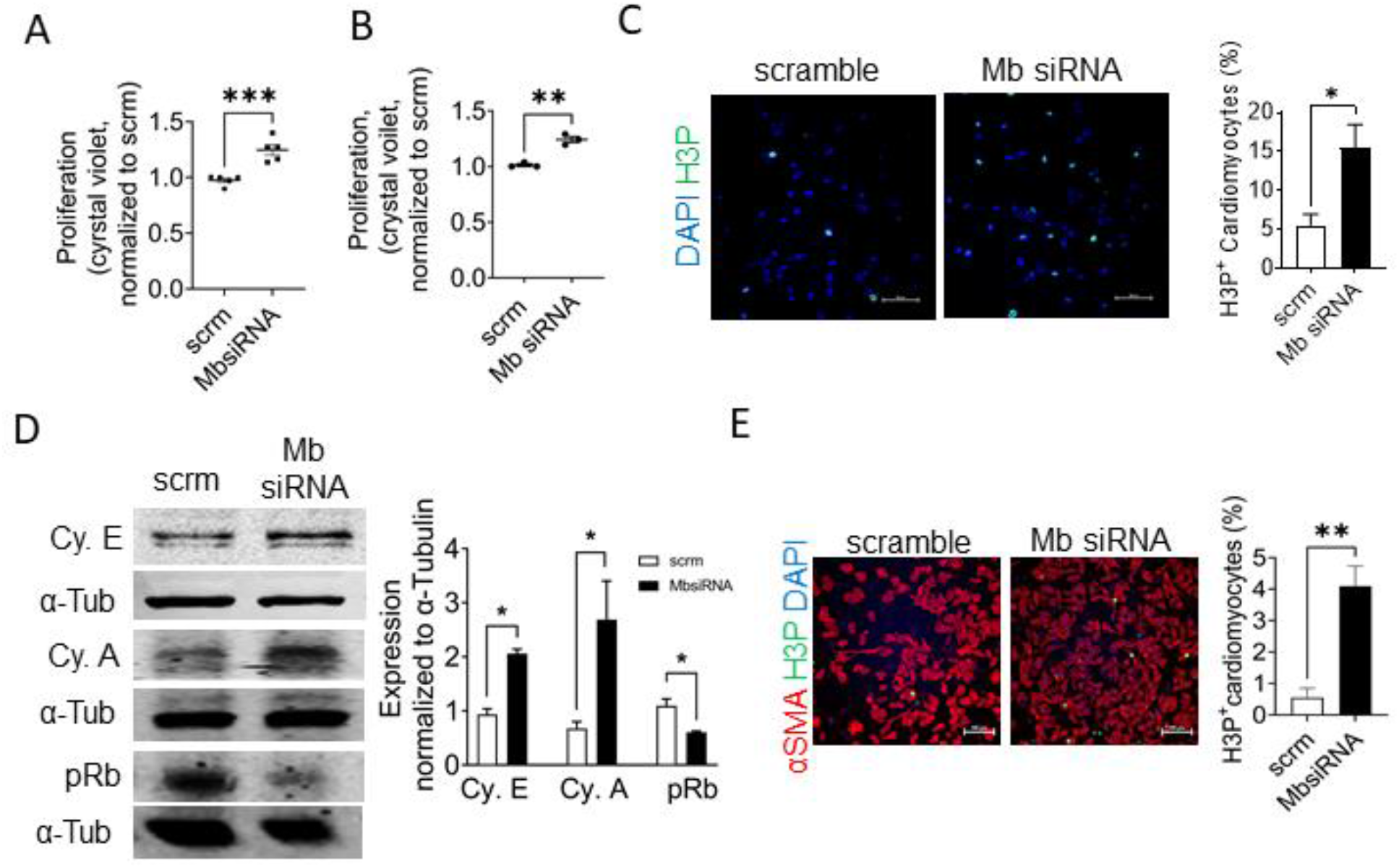
Knockdown of Mb stimulates cardiomyocyte proliferation. **(A)** Quantification of cell proliferation in H9C2 cells expressing scrm and Mb siRNA Mb cells, expressed as fold change of scramble control; n=5. **(B)** Quantification of cell proliferation in HL-1 cells expressing scrm and Mb-siRNA by crystal violet incorporation; n=3. **(C)** Representative immunofluorescence images and quantification of phospho-Histone3 (H3P) positive cells in M-phase (green) and nuclei (blue) in scrm control and Mb siRNA H9C2 cells. Quantification is from 6 high power fields/sample; n=3. **(D)** Representative Western blots and the corresponding quantifications measuring Cyclin E, A (Cy.E&A) and phospho-Retinoblastoma (pRb) in scrm and Mb siRNA H9C2 cells; n=3. **(E)** Representative immunofluorescence images and quantification of phospho-Histone3 (H3P) positive cells in M-phase (green) and nuclei (blue) in scrm control and Mb siRNA NRVMs; n=3. \**p*< 0.05, \*\**p*< 0.01, \*\*\**p<0*.*001*. Bar graphs depict mean ± SEM.

### Myoglobin promotes differentiation by YAP inactivation

We next sought to determine whether Mb promotes differentiation through the modulation of the transcriptional co-activator YAP. Phosphorylation of YAP maintains the co-activator in an inactive state, while dephosphorylation activates YAP, enabling its nuclear translocation and transcriptional activity to drive cardiomyocyte proliferation and inhibit differentiation^33, 48^. Knockdown of Mb in H9C2 cells significantly decreased the levels of YAP phosphorylation (pYAP) at Ser127, a key phospho-site responsible for its inactivation, without changing total YAP (T-YAP) levels (**Figure 3A**). Mb-siRNA cells also demonstrated greater nuclear localization of YAP compared to scrm controls (**Figure 3B**). To test whether Mb-dependent phosphorylation altered YAP transcriptional activity, we transfected scrm and Mb-siRNA cells with a luciferase reporter to measure the transcription of TEAD, a downstream target of YAP. Consistent with YAP activation in cells lacking Mb, MbsiRNA cells showed increased transcription of TEAD, confirming Mb knockdown increases YAP transcriptional activity (**Figure 3C**). Measurement of the mRNA levels of two classical gene targets of YAP, *ctgf* and *cyr61*, also showed higher levels in Mb-siRNA cells (**Figure 3D**), consistent with the TEAD reporter assay. Notably, compared to overexpression of a wildtype Mb plasmid, overexpression of a Mb mutant lacking the heme moiety (Apo-Mb) did not increase the levels of pYAP in H9C2 cells lacking endogenous Mb (**Figure 3E**). Collectively, these results demonstrate that the loss of Mb decreases YAP phosphorylation, and enables its translocation to the nucleus, which results in a potentiation of its transcriptional activity. Further, this effect is dependent on the heme center of Mb.

**Figure 3:**
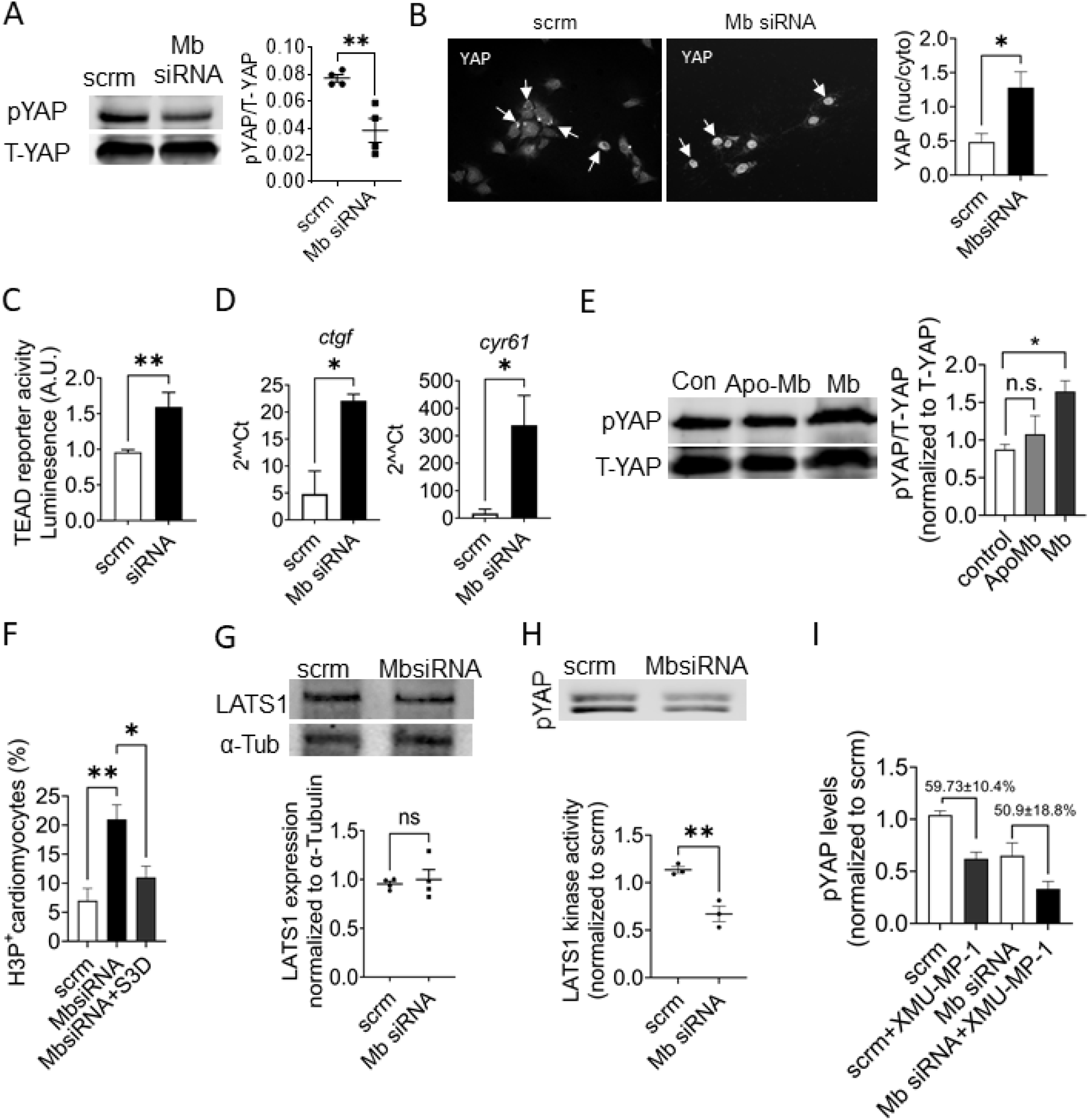
Myoglobin propagates YAP phosphorylation and LATS1 kinase activity. **(A)** Representative Western Blot and quantification of phospho-YAP (pYAP) and total YAP (T-YAP) in scrm and Mb-siRNA cells 48 hours after siRNA transfection; n=4. **(B)** Representative immunofluorescence images depicting YAP cellular localization in scrm and Mb-siRNA cells. Arrows point to instance of cytoplasmic signal (scrm image) and nuclear signal (MbsiRNA image). Quantification is the ratio of nuclear to cytoplasmic staining intensity of YAP from 6 high-power field per sample; n=3. **(C)** TEAD-luciferase reporter signal in scrm controls, Mb-siRNA cells; n=5. **(D)** RNA levels of YAP transcriptional target genes -*ctgf* and *cyr61*-in scrm and Mb-siRNA cells; n=3. **(E)** Representative Western blot for pYAP and T-YAP levels in a stable cell line with Mb knockdown (control; con), and overexpression of Mb or Apo-Mb for 48 hours. n=3. **(F)** Quantification of M-phase cell cycle activity in scrm, Mb siRNA and Mb siRNA cells overexpressing YAP phosphomimetic S3D; n=4. **(G)** Representative Western blot and quantification of LATS1 protein level in scrm and Mb-siRNA cells. n=4. **(H)** Kinase activity of immunoprecipitated LATS1 from scrm and Mb-siRNA cells; n=3. **(I)** pYAP levels following XMU-MP-1 treatment in scrm control and MbsiRNA cells. n=3. \**p*< 0.05, \*\**p*< 0.01, ns= not significant. Bar graphs depict mean ± SEM.

To test whether decreased YAP phosphorylation was required for the undifferentiated/proliferative phenotype observed with the deletion of Mb, we utilized a phosphomimetic YAP mutant, S3D, which remains constitutively phosphorylated^49^. While MbsiRNA cells had increased phospho-histone 3 levels indicating more proliferative activity as expected, overexpression of YAP S3D in Mb-siRNA cells inhibited this increase (**Figure 3F**), demonstrating that Mb regulates cellular differentiation through the modulation of YAP phosphorylation.

### Mb-derived ROS potentiates LATS1 kinase activity

To elucidate the mechanism by which Mb regulates YAP phosphorylation, we focused on LATS1/2, the upstream Hippo pathway kinase that phosphorylates YAP. LATS1 protein levels were not different between scrm and MbsiRNA cells (**Figure 3G**). However, the kinase activity (the ability to phosphorylate YAP) of LATS1 immunoprecipitated from Mb-siRNA cells was significantly lower than compared to LATS1 isolated from control cells (**Figure 3H**). These data suggest that Mb supports LATS1 kinase activity to increase YAP phosphorylation.

In the canonical Hippo pathway, LATS1 activity is regulated by the upstream kinase MST1/2. To test whether differential MST1/2 kinase activity was responsible for the difference in LATS1 activity between MbsiRNA and control cells, we measured pYAP levels from scrm and Mb-siRNA cells treated with a pharmacological inhibitor of MST1/2 (XMU-MP-1; 10µM, 1h).^50^ Inhibition of MST1/2 decreased pYAP levels in scrm and MbsiRNA cells to the same extent (59.73 ± 10.4% reduction in scrm cells vs 50.9 ± 18.8% reduction in Mb siRNA cells) (**Fig 3I**). These data are consistent with the idea that in Mb-expressing cells, LATS1 activity is regulated by an additional mechanism independent of MST1/2.

It is well established that kinase activity can be regulated by reactive oxygen species (ROS) and the heme moiety of Mb can catalyze the production of ROS^16, 51^. To test whether Mb-dependent ROS production modulates LATS1/YAP signaling, we first assessed cellular ROS production by scrm control and Mb-siRNA cells. In accordance with prior studies^51^, MbsiRNA cells generated lower levels of total cellular ROS than scrm cells (**Figure 4A**) while no difference was observed in the levels of mitochondrial-derived ROS (**Figure 4B**). Furthermore, while overexpression of wildtype Mb in H9C2 cells lacking endogenous Mb increased the cellular ROS levels, overexpression of Apo-Mb did not increase ROS production, demonstrating that Mb mediated ROS production is heme-dependent (**Figure 4C**).

**Figure 4:**
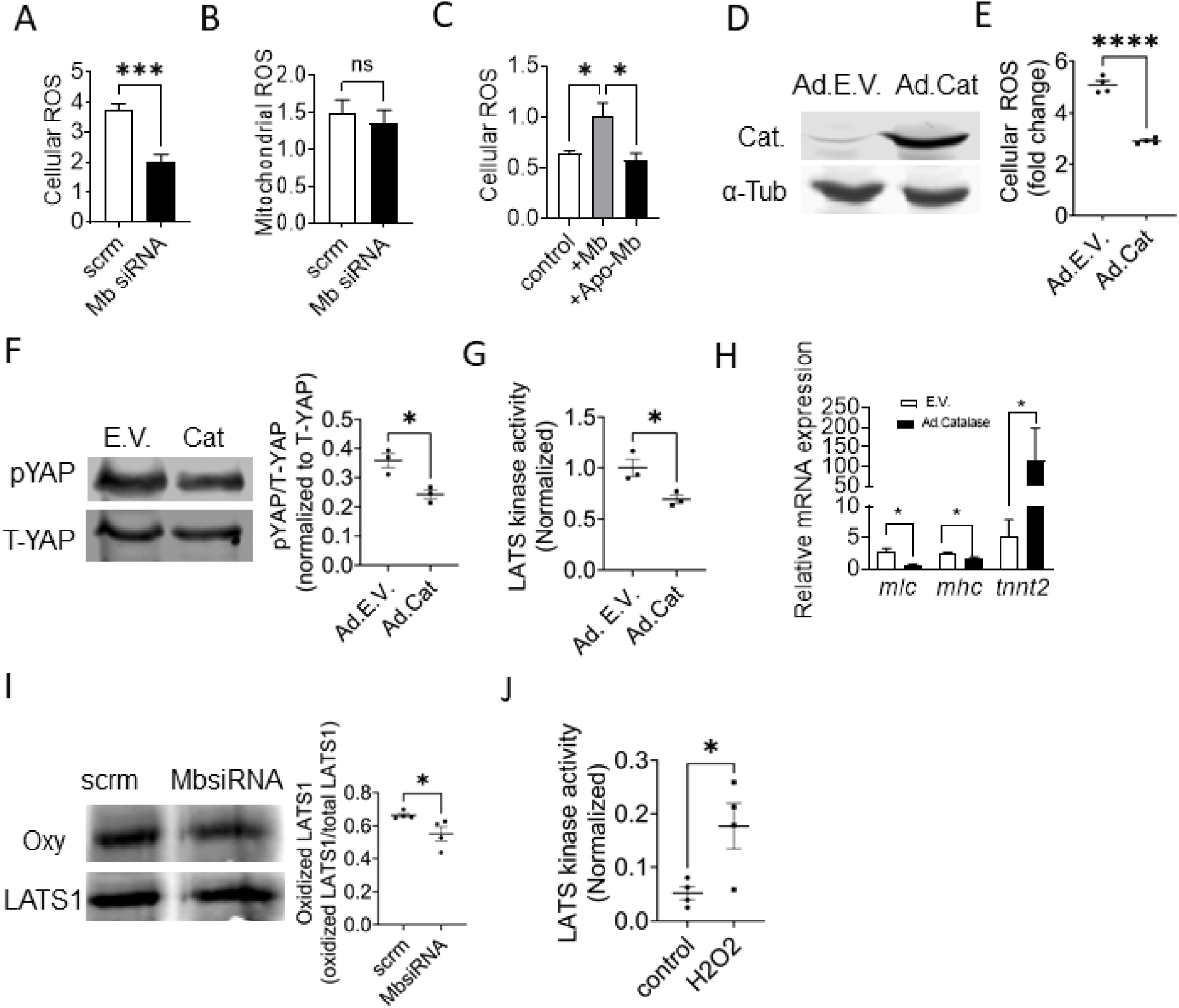
Scavenging ROS inhibits whereas oxidation increases LATS1 activity. **(A)** Cellular and **(B)** mitochondrial ROS levels in scrm and Mb-siRNA cells; n=4-6. **(C)** Cellular ROS levels in H9C2 cells with Mb knockdown: control, Mb overexpression and Apo-Mb overexpression; n=3. (**D**) Representative western blot for catalase expression in H9C2 cells transduced with empty vector or adenoviral catalase transduction. **(E)** Cellular ROS levels in H9C2 cells transduced with empty vector (Ad EV) or adenoviral catalase (Ad Cat); n=4. **(F)** Representative Western blot and quantification for pYAP and total YAP levels in H9C2 cells transduced with empty vector (Ad EV) or with adenoviral catalase (Ad Cat) expression; n=3. **(G)** Kinase activity of LATS1 isolated from empty vector (Ad EV) or adenoviral catalase (Ad Cat) transduced H9C2 cells; n=3. **(H)** RNA levels of cardiac differentiation markers in cells overexpressing empty vector control and catalase adenovirus; n=3. **(I)** Levels of oxidized LATS1 kinase immunoprecipitated from scrm and Mb-siRNA cells by oxyblot assay. n=4. **(J)** LATS1 kinase activity with and without treatment with hydrogen peroxide (100 µM; 2 min); n=3. \**p* <0.05, \*\*\**p*<0.001. Bar graphs depict mean ± SEM.

To directly test whether Mb-dependent ROS regulates LATS1 activity, H9C2 cells were transduced with adenovirus to overexpress catalase (**Figure 4D**, inset), which catalyzes the breakdown of hydrogen peroxide. Compared to empty vector control, catalase overexpression decreased cellular ROS levels as expected (**Figure 4E**). Notably, catalase overexpression had little effect on cellular ROS levels in MbsiRNA cells. Catalase overexpression in control cells also significantly decreased pYAP levels in Mb containing cells, (**Figure 4F**) and decreased the kinase activity of LATS1 (**Figure 4G**), mimicking the effect of Mb knockdown. Similar to Mb-siRNA cells, overexpression of catalase in scrm cells also inhibited the expression of the differentiation markers *mlc, mhc* and *tnnt2* (**Figure 4H**).

Based on the observation that cellular ROS potentiated LATS1 activity, we tested whether LATS1 protein could be oxidatively modified. Utilizing an oxyblot assay, we measured Mb-dependent differential oxidative modifications of cellular proteins. Myoglobin-expressing scrm cells showed a higher level of oxidized LATS1 kinase compared LATS1 kinase in Mb siRNA cells (**Figure 4I**). We next tested whether oxidative modification could impact LATS1 activity. Exposure of purified LATS1 active peptide to hydrogen peroxide for 5 minutes *in vitro* significantly increased its kinase activity (**Figure 4J**). Collectively, these results demonstrate that Mb-dependent oxidation of LATS1 increases its kinase activity.

### Loss of Mb drives a pro-proliferative and regenerative response in zebrafish after cardiac injury

In contrast to mammals, zebrafish (*Danio rerio*) can regenerate damaged myocardium throughout their lifetime. Zebrafish heart regeneration occurs via multiple coordinated events^10, 11, 15^. A key feature is the extensive cardiomyocyte dedifferentiation and proliferation post-injury which primarily contributes to regeneration of damaged myocardium and long-term resolution of scar tissue. To determine whether Mb regulates YAP activity and cardiomyocyte proliferation *in vivo*, we generated mutant zebrafish deficient in Mb (*mb-/-*; **Figure 5A**). We confirmed loss of Mb expression by immunofluorescence (**Figure 5B**) and Western blot (**Figure 5C**). Consistent with our in vitro data, Mb deletion decreased the levels of pYAP in the zebrafish heart (**Figure 5C**). In zebrafish hearts, differentiated cardiomyocytes are able to regress to a de-differentiated state and subsequently proliferate in response to injury^10^. We employed a model of myocardial amputation in the zebrafish to test whether endogenous Mb regulates cardiomyocyte proliferation and heart regeneration. At 7 days following ventricular amputation, Mb-/-hearts showed a significantly higher number of dedifferentiated cardiomyocytes identified by staining for embryonic cardiac myosin heavy chain (embCMHC) (**Figure 5D**) and a higher number of proliferating cardiomyocytes compared to WT hearts– proliferation index 0.219 ± 0.057 in WT vs .391 ± .128 in *mb-/-*(**Figure 5E**)^52^. To determine whether loss of Mb impacts long-term regeneration, we examined hearts for scar resolution using AFOG staining. Zebrafish lacking Mb had significantly smaller scar area (4.07 ± 1.74% area) at 20 days post amputation (dpa) compared to WT hearts (10.12 ± 5.92% area). Notably, Mb-/- zebrafish hearts showed no significant change in scar sizes between 20 and 30 dpa, likely due to the fact that scarring was predominantly resolved at 20 dpa (**Figure 5F**)^10^.

**Fig 5:**
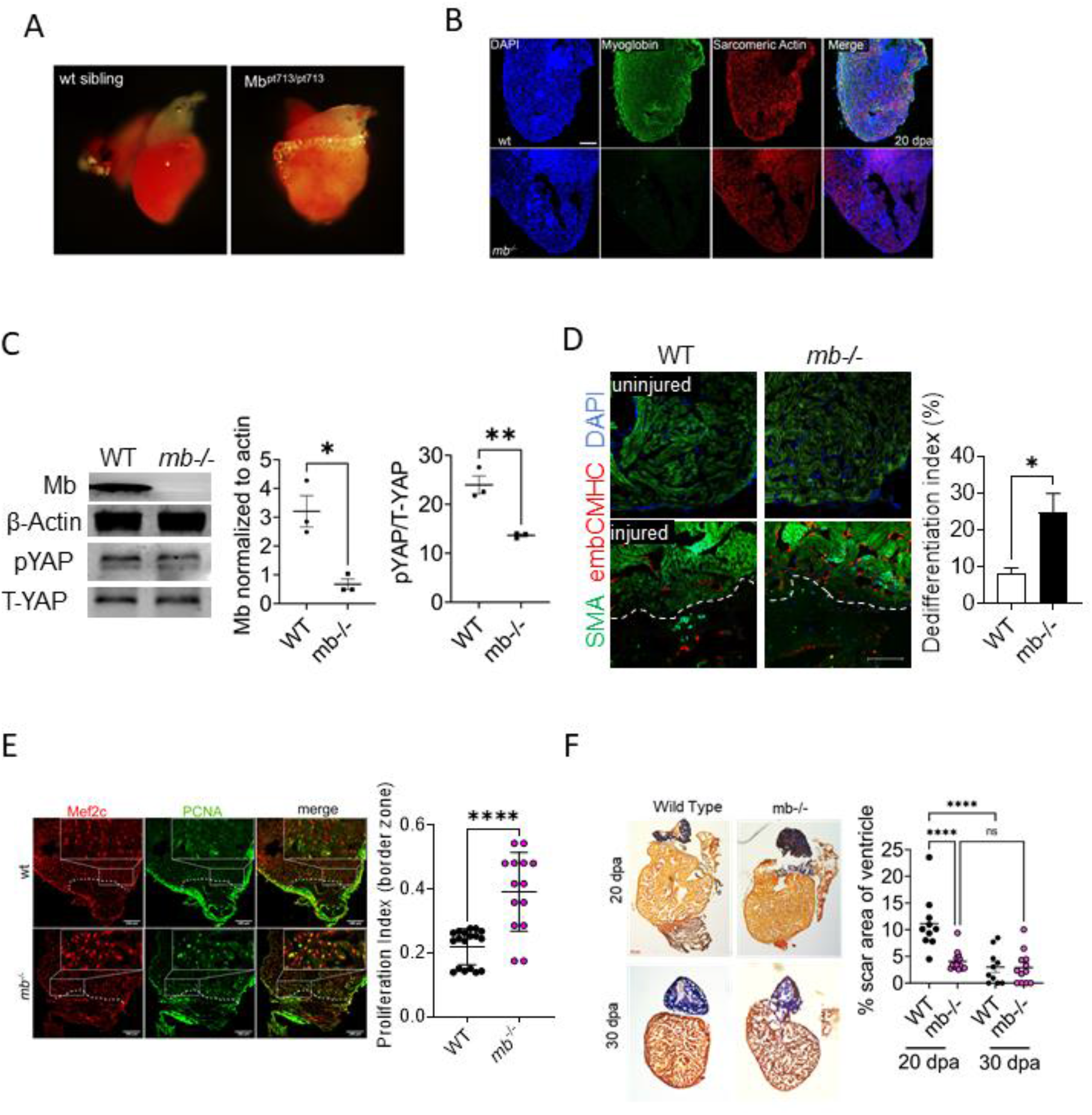
Loss of Mb promotes cardiomyocyte proliferation and regeneration *in vivo*. **(A)** Representative images of whole zebrafish hearts from fish expressing (WT) or lacking Mb (mb -/-). **(B)** Immunofluorescence images of whole heart sections from wt and *mb-/-* zebrafish stained with DAPI (blue) to visualize nuclei, Mb protein (green), sarcomeric actin (red) and their co-localization. **(C)** Representative Western blot and quantification of Mb, pYAP and total YAP levels in WT and *mb-/-* zebrafish hearts; n=3. **(D)** Zebrafish heart section demonstrating the staining for dedifferentiation marker embCMHC in red and α-sarcomeric actin in green at 7 days post amputation, and quantification of embCMHC cardiomyocytes (Differentiation index: % of cardiomyocytes expressing embCMHC in border zone of injury) from WT and *mb-/-* hearts; n=6. (**E**) Heart sections from WT and mb-/- zebrafish stained for Mef2c transcription factor (red) to identify cardiomyocytes and PCNA (green) to identify proliferating cells at 7 days post amputation; n=7-9. **(F)** Representative histological sections of AFOG stained tissue from WT and *mb-/-* zebrafish at 20- and 30-days post amputation stained to visualize fibrin (red), collagen (blue) and cardiac muscle (orange). Graph demonstrates the quantification of scar area on each section normalized to total ventricular area; n=9-16 *\*p<0*.*05*, \*\**p*<0.01, \*\*\*\**p*<0.0001. Bar graphs depict mean ± SEM.

## Discussion

The major finding of this study is that endogenous Mb expression propagates and maintains cardiomyocyte differentiation, and mechanistically, this is dependent on Mb-mediated oxidative regulation of the Hippo signaling pathway. Specifically, we show that Mb-dependent oxidation of LATS1 kinase potentiates its activity to phosphorylate and inactivate YAP, which inhibits its downstream transcriptional activity. Further, we show that this pathway is active *in vivo* as zebrafish lacking Mb show accelerated cardiac regeneration after injury. This study delineates a novel physiological role for Mb that has implications not only for cardiomyocyte maturation in cardiac development but also for potential regenerative therapeutic applications.

Our data suggest a fundamental physiological role for Mb in propagating and maintaining cardiomyocyte differentiation, distinct from its traditional function as an oxygen storage and delivery protein. While the role of Mb as a catalyst of oxidative stress has been examined in pathological contexts such myocardial infarction,^22^ a physiological role for its oxidative signaling in cardiomyocytes has not been defined. Our data are consistent with an earlier report of hydrogen peroxide treatment inducing H9C2 cell cycle arrest prior to differentiation, though the effect of this exogenous oxidant is more nuanced depending on timing and dose of exposure.^53^ Importantly, the role of oxidants in cardiomyocyte maturation is established. Cellular ROS have been implicated in the mechanical or electrical stimuli-induced differentiation of embryonic cells to cardiomyocytes.^54-56^ In contrast, scavenging or decreasing the levels of ROS using pyrrolidine dithiocarbamate, catalase, or N-acetylcysteine have been shown in other models to inhibit the formation of cardiomyocytes from developing embryoid bodies.^57^ Despite their significance in cardiomyocyte differentiation, the endogenous source of ROS responsible for potentiation of differentiation remains less clear. Prior research has largely focused on the role of NADPH oxidases (NOX) enzymes as a source of intracellular oxidants in stimulating cardiomyocyte differentiation. Downregulation of NOX4 during early stages of cardio myogenesis inhibited cardiomyocyte differentiation.^56, 57^ The role of other sources of ROS are less clear. For example, conflicting studies demonstrate that ROS derived from mitochondria either stimulates or inhibits cardiomyocyte differentiation^58, 59^ and can also either induce cardiomyocyte cell cycle arrest^60, 61^ or promote proliferation in the context of T3 signaling.^62, 63^ Our results demonstrate Mb as a major source of cardiomyocyte ROS which significantly impacts cardiomyocyte differentiation It is established that the levels of Mb dynamically increase during development from the fetal to adult stage, and that this correlates with the development time period of increased cardiomyocyte differention.^24^ However, how this change alters cardiac development is unknown. Studies from the global knockout mouse demonstrate that Mb is critical for cardiac development as the high proportion of global Mb knockout mice that fail to develop physiological compensatory mechanisms die embryonically.^28^ However, further study is required to link the loss of Mb expression to decreased oxidative signaling and deficient differentiation in these mice.

Despite its significance in development, homeostasis and tissue repair, the endogenous mechanisms regulating the Hippo pathway and the phosphorylation of YAP in cardiomyocytes are yet to be completely uncovered^64^. The data herein demonstrate that Mb-mediated oxidative signaling is a critical regulator of the Hippo pathway that maintains YAP in its phosphorylated state. Prior studies utilizing exogenous oxidants, such as hydrogen peroxide have shown that the upstream kinase MST1 can be oxidatively modified and activated in a Prdx1 dependent manner. ^65-67^ While it is possible that Mb-dependent ROS can alter MST1 to alter LATS1 activity, our data with the MST1 inhibitor (XMU-MP-1) as well as with the purified LATS1 peptide suggests an additional mechanism of LATS1 regulation, independent of MST1. Our study thus extends the paradigm of oxidative activation of the Hippo kinase cascade from the MST1 kinase to LATS1 kinase. The identification of Mb as a regulator of YAP phosphorylation not only demonstrates an endogenous oxidative mechanism of regulation of Hippo signaling, but potentially links Hippo signaling to the oxygen sensing properties of Mb, enabling a more intricate Mb-dependent signaling axis in the regulation of development and tissue repair.

In cardiomyocytes, LATS1 and LATS2 have overlapping functions in phosphorylating YAP^34^, and while our studies are currently limited to LATS1 kinase, it does not exclude LATS2 modification in a similar mechanism. Future studies will examine the role of cellular oxidants on LATS1/2 activity across different cell types, given the conserved nature of the pathway in multiple organs.^34, 68, 69^ Beyond cellular homeostasis, active Hippo signaling functions as a tumor suppression mechanism in multiple cell types, and dysregulation of this pathway or hyperactivation of its effector YAP are extensively associated with cancer progression. We and others have reported that certain epithelial cancers aberrantly express Mb.^51^ Our results also demonstrated that expression of holo-Mb but not Apo-Mb significantly inhibited proliferation of breast epithelial cancer cells, confirming a role for Mb-derived oxidants in regulating cancer cell fate. Examining the role of Mb on Hippo signaling beyond cardiomyocytes has the potential to advance the field of tissue growth and repair in other areas, such as in the development of anti-cancer therapies.^70^

Elucidation of the Mb-LATS1-YAP pathway provides a roadmap to potential therapeutics for cardiac repair. *In vivo* apical resection of the heart in our zebrafish model provides proof-of-concept that that Mb deletion drives cardiomyocyte de-differentiation and proliferation. \. While a limitation of these studies is the use of zebrafish and further study is required to test the pathway in mammalian models and humans, these observations suggest that loss of Mb may be protective following cardiac injury. One study that potentially provides evidence to support this hypothesis is that in which global Mb knockout mice infused with isoproterenol for 14 days were protected from LV dilation and development of heart failure compared to wildtype mice which expressed Mb.^27^ Indeed, it is well-documented that following injuries such as myocardial infarction or in progressive cardiomyopathies, there is a downregulation of endogenous Mb expression in multiple species including humans.^71-73^ However, whether this downregulation modifies the course of recovery has not been considered and our results bring this question to the forefront of the field. Beyond development and endogenous repair, our data also offer a new strategy to contribute to the field induced pluripotent cell (iPSC) technology, which offers a viable source of cardiomyocytes to replace damaged tissue and integrate with the spared myocardium. It is possible that modulation of Mb-dependent cardiomyocyte differentiation can be translated to optimize protocols for increasing efficiency of iPSCs.^74-76^ While our data shows that Mb can drive regenerative events via YAP^32-35, 76-79^, future studies should focus on whether Mb can affect reparative processes beyond these pathways. The possible mechanisms include altered oxidative stress response,^80^ cellular metabolism^81, 82^, promoting expression of other fetal genes^75, 81, 83, 84^, or altering the extracellular environment.^48, 85-88^It is also interesting to speculate that Mb, through its more recently described chemistry with reactive nitrogen species or fatty acid binding^17, 89-98^, could impact the functional properties of cardiomyocytes to impact regeneration.

In summary, we have described a novel pathway by which Mb regulates cardiomyocyte differentiation. We show that this occurs via Mb-dependent ROS which regulates YAP/LATS1 signaling. The loss of Mb is beneficial for cardiac regeneration, in part through promoting substantial cardiomyocyte dedifferentiation. Taken together, these results have strong implications for the role of Mb in regulating Hippo signaling, and for advancing the understanding of cardiac function under physiological and pathological conditions.

## List of non-standard abbreviations

Mb: Myoglobin
NRVM: neonatal rat ventricular myocytes
YAP: Yes associated protein
LATS: Large tumor suppressor kinase
ROS: Reactive Oxygen species
Dpa: Days post amputation
H3P: Phospho-histone H3
Mef2c: Myocycte enhancer factor 2c

## Acknowledgements

We would like to thank Jianmin Xu for her technical expertise contributed towards this work. We would like to thank Dr. Jixin Dong, Eppley Institute for Research in Cancer and Allied Diseases, University of Nebraska Medical Center for kindly sharing the YAP SD mutant with us. The N2.261 embCMHC antibody developed by Dr David R Soll, was obtained from the Developmental Studies Hybridoma Bank, created by the NICHD of the NIH and maintained at The University of Iowa, Department of Biology, Iowa City, IA 52242.

